## Supplementary material for "Glycolytic pyruvate kinase moonlighting activities in DNA replication initiation and elongation": Fig S1-S8

**A** Key residues of Cat domains

| PykA *B. subtilis* | Human PKM2 | PYK *Mycobacterium tuberculosis* |
|---|---|---|
| R32 | R73 | R33 |
| R73 | R120 | R74 |
| K220 | K270 | K218 |
| G245 | G295 | G243 |
| D246 | D296 | D244 |
| T278 | T328 | T276 |

**B** Alignment of PEPut domains

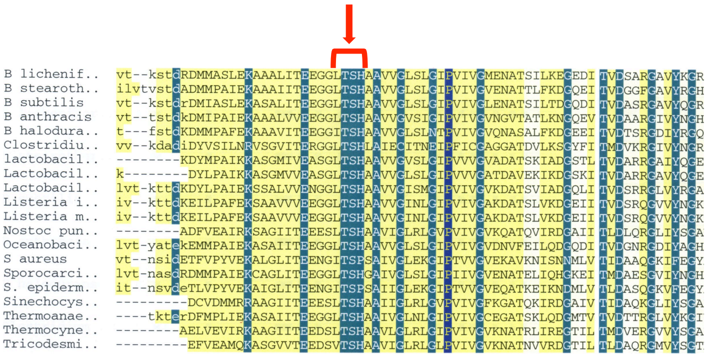

**Fig. S1**: Key amino-acids of the Cat and PEPut domain of PykA.

**A.** Cat domain analysis. Clustalw and Chimera analysis of the pyruvate kinase of *B. subtilis* (PykA), human cells (PKM2), *Mycobacterium tuberculosis* (PYK) and *Geobacillus stearothermophilus* identified key amino acids of the catalytic site of the *B. subtilis* PykA protein.

**B.** PEPut domain analysis. Alignment of the PEPut domain of PykA to related domains of various metabolic enzymes. The red arrow highlights the conserved LTSH motif.

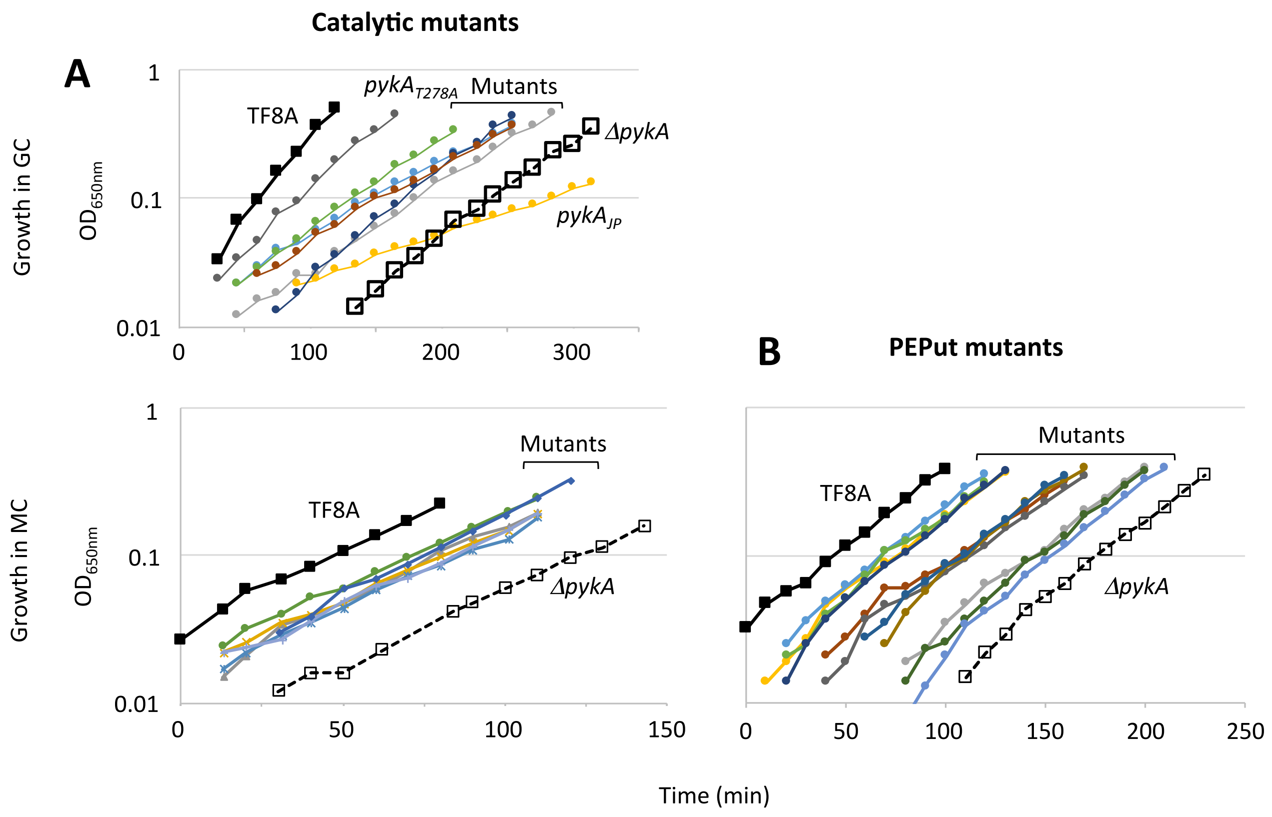


**Fig. S2**: Effect of Cat and PEPut mutations on growth in the GC and MC media.

Wild-type and *pykA* mutants were first grown over-night in the MC or GC medium supplemented with appropriate antibiotic. Upon saturation, cultures were diluted 1000-fold in the same medium without antibiotic and growth was monitored spectrophotometrically.

**A.** Analysis of catalytic mutants ($pykA_{\Delta cat}$, $pykA_{R32A}$, $pykA_{R73A}$, $pykA_{K220A}$, $pykA_{GD245/6AA}$, $pykA_{T278A}$, $pykA_{JP}$). Controls: TF8A (wild-type) and *ΔpykA*.

**B.** Analysis of PEPut and Cat-PEPut interaction mutants ($pykA_{\Delta PEP}$, $pykA_{T>A}$, $pykA_{S>A}$, $pykA_{H>A}$, $pykA_{TSH>AAA}$, $pykA_{T>D}$, $pykA_{S>D}$, $pykA_{H>D}$, $pykA_{TSH>DDD}$, $pykA_{E209A}$, $pykA_{L536A}$). Controls: TF8A (wild-type) and *ΔpykA*.

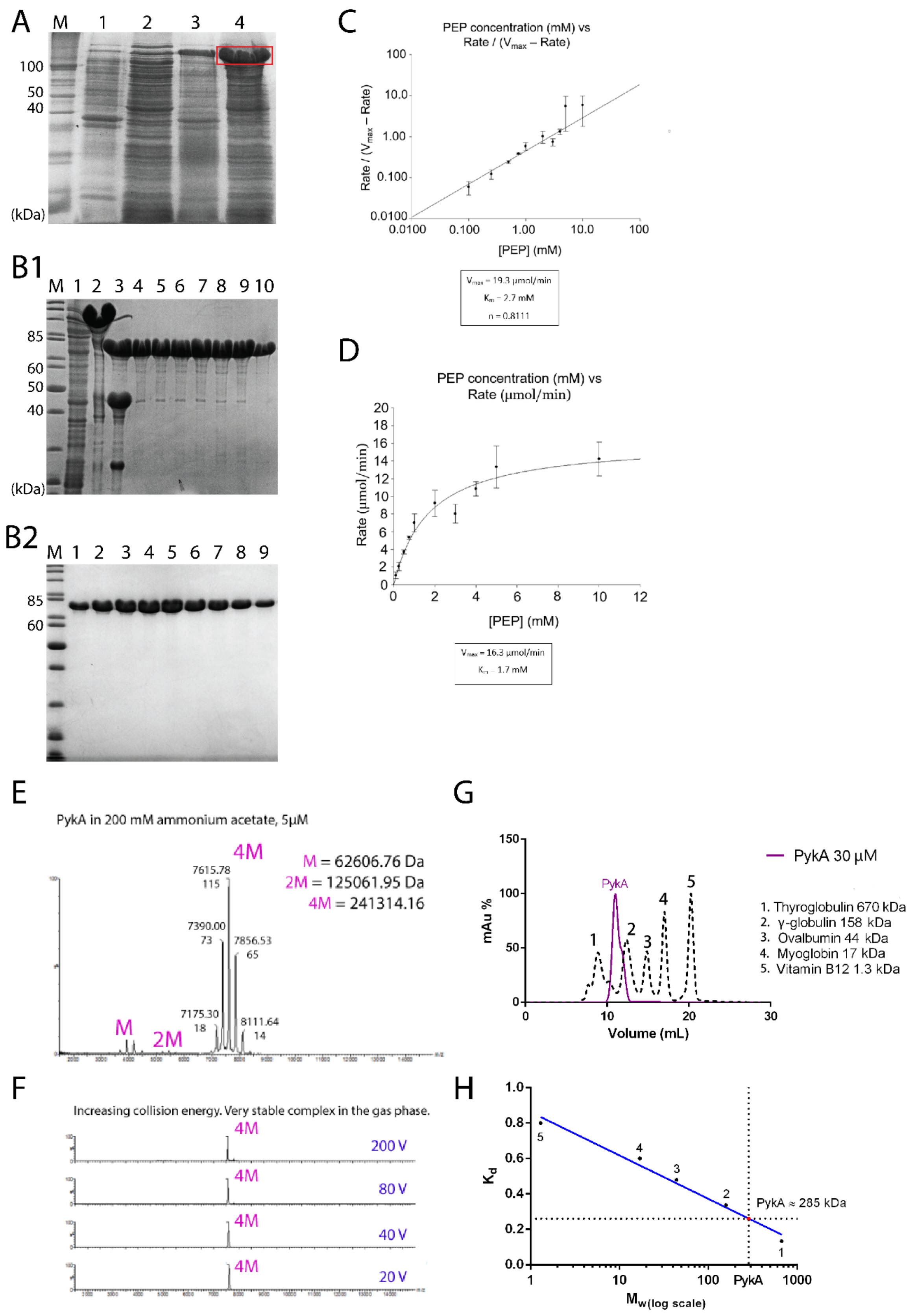


**Fig. S3:** PykA purification and characterization of its function and oligomeric state.

**A.** SDS-PAGE (15% polyacrylamide gel) showing over-expression of the 6His-MBP-PykA in Rosetta (DE3) *E. coli.* The soluble expressed tagged PykA protein is shown in a red rectangular in lane 4, whereas lanes M, 1, 2 and 3 show molecular weight standards, the control uninduced insoluble fraction, the control uninduced soluble fraction and the IPTG-induced insoluble fraction, respectively.

**B.1.** SDS-PAGE (15% polyacrylamide gel) showing fractions from the first IMAC purification step of PykA. From left to right, lanes represent molecular weight standards (M), the flow through (1), the eluted tagged PykA (2), the overnight TEV treated tagged PykA (3), the flow through fractions containing untagged PykA from the second IMAC step after TEV proteolysis (4-10). **B.2.** SDS-PAGE (15% polyacrylamide gel) showing the final gel filtration column (HiLoad 26/60 Superdex 200 Prep Grade Gel Filtration Column). From left to right, lanes represent molecular weight standards (M) and fractions of the size exclusion chromatography (1-9).

**C.** The graph shows a Hill plot for the activity of PykA at 25$^{o}$C. The Rate/(Vmax-Rate) (Y-axis) was plotted against the PEP substrate concentration (X-axis) using GraphPad Prism 4 software and the Vmax (19.3 µmol/min), Km (2.7 mM) and the Hill coefficient n (0.8111) values are shown below the graph. The n value is <1 indicating negative cooperative binding of PykA to its PEP substrate.

**D.** The graph shows a Michaelis-Menten plot for the activity of PykA at 25$^{o}$C. The initial rate of the reaction (Y-axis) was plotted against the PEP substrate concentration (X-axis) using GraphPad Prism 4 software and the Vmax (16.3 µmol/min) and Km (1.7 mM) values are shown below the graph.

**E.** A native mass spectrum showing the PykA tetramer and miniscule amounts of the dimer and monomer. The theoretical mass of the PykA monomer (62,314.9 Da), dimer (124,629.8 Da) and tetramer (249,259.6 Da) match very well with the calculated masses of the monomer (62,606.8 Da), dimer (125,062 Da) and tetramer (241,314.2 Da) from the spectrum.

**F.** Increasing the collision energy gradually from 20, 40, 80 to 200 V did not affect the PykA tetramer indicating a very strong tetramer in the gas phase.

**G.** Comparative analytical gel filtration of the PykA tetramer against molecular weight standards (Thyroglobulin 670 kDa, g-globulin 158 kDa, ovalbumin 44 kDa, myoglobin 17 kDa and vitamin B12 1.3 kDa) through a Superdex 200 10/300 GL prepacked Tricorn gel filtration column (GE Healthcare).

**H.** Selectivity trendline made from the molecular weight standards (shown in graph G) for the estimation of the PykA MW. The x axis is in logarithmic scale. Graphpad was used for plotting the data points. The theoretical value of our PykA (249,259.6 Da) is close to the estimated (285,000 Da) which along with the MS data verifies the tetramer in solution. $K_d$ is the equilibrium distribution coefficient. The numbers (1-5) on the data points correspond to the proteins shown in graph G.

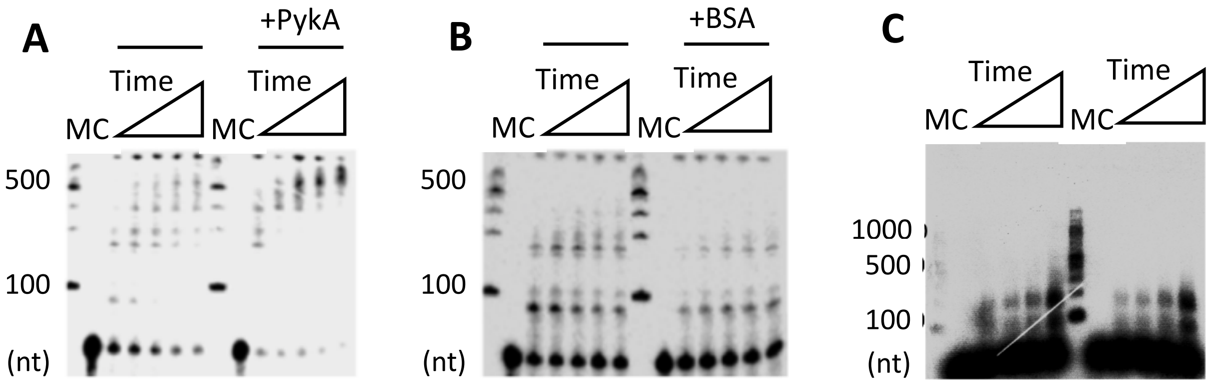


**Fig. S4**: Stimulation of DnaE activity by PykA but not by BSA.

**A.** Primer extension assays monitoring the extension of a 5'-$^{32}$P-radioactively labelled 60mer DNA primer annealed onto M13 ssDNA over time by the *B. subtilis* DnaE. The activity of DnaE polymerase (10 nM) was monitored in the presence and absence of PykA (10 nM, tetramer) through a time course (30-150 sec). Lanes in the gels from left to right indicate: (M): DNA-ladder and then the time course (0, 30, 60, 90, 120 and 150 sec) depicted by the rectangular triangle.

**B.** Primer extension assays as above with or without 10 nM (monomer) BSA instead of PykA.

**C.** DnaE (1 nM) polymerase activity at increasing BSA concentrations (0, 5, 50, 500 nM), as indicated by the rectangular triangle, monitored by alkaline agarose electrophoresis. The DNA substrate is a labelled 20mer (5'-CAGTGCCAAGCTTGCATGCC-3') primer annealed onto ssM13 ssDNA (2nM). The primer extension reaction was carried out for a longer time than above (5 min instead of 30-150 sec) and the film was over-exposed to compensate for the lower DnaE concentration. The assay was carried out at 37 °C in 50 mM Tris-HCl 7.5, 50 mM NaCl, 10 mM $MgCl_2$ mM DTT, 1 mM dNTPs. No stimulation of the DnaE polymerase activity was observed in the presence of 5 and 50 nM BSA. The marginal stimulation observed at 500 nM BSA excess is likely because at this high concentration, BSA acts as a blocking agent preventing adhesion of DnaE to the plastic reaction tubes.

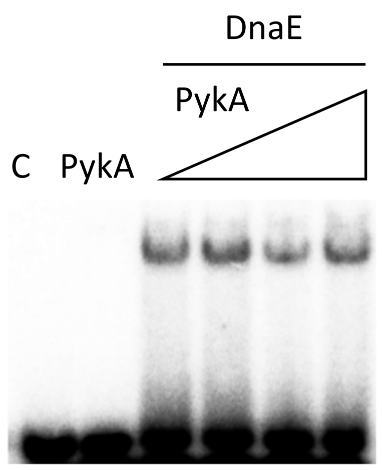


**Fig. S5:** Stimulation of DnaE activity by PykA does not result from stimulation of DnaE binding to primed templates.

EMSA investigation of the effect of PykA on the DNA binding of DnaE polymerase. The DNA substrate was constructed by annealing a 5'-$^{32}$P-radioactively labelled 15mer (5'-AAGGGGGTGTGTGTG-3') primer annealed onto a 30mer (5'-ACACACACACACACACACACACACCCCCTT-3') oligonucleotide. Binding reactions were carried out with 1 nM DNA substrate, DnaE (500nM) and increasing concentrations (0, 12.5, 125 and 1,250 nM tetramer) of PykA, as indicated by the rectangular triangle for 10 min at 37 °C in 50 mM NaCl, 10 mM $MgCl_2$, 50 mM Tris-HCl pH 7.5. Lanes C and PykA represent the radioactive substrate in the absence of any proteins and in the presence of PykA (1,250 nM tetramer), respectively, showing that PykA does not bind to the DNA substrate. No stimulation of DnaE binding to DNA was observed in the presence of increasing concentrations of PykA indicating that PykA does not enhance the DNA binding activity of DnaE.

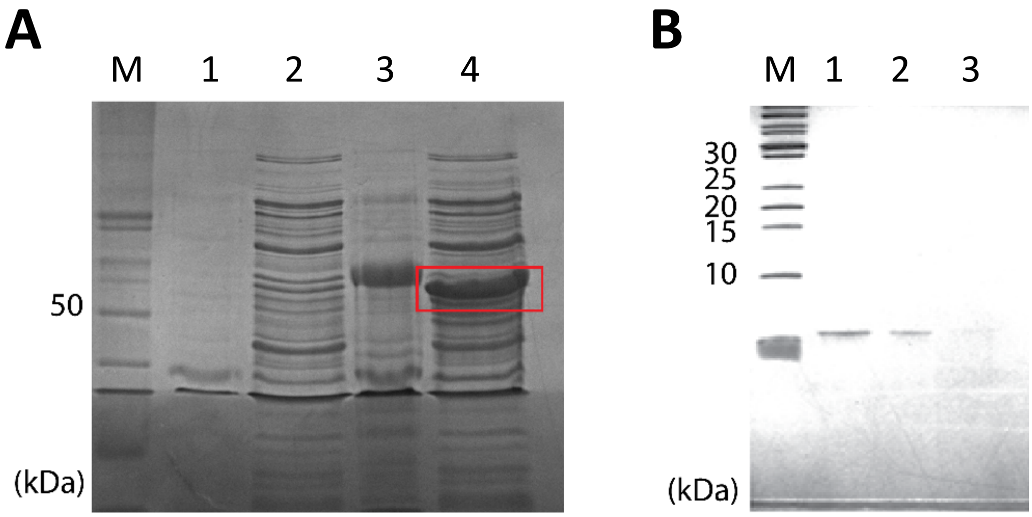


**Fig. S6:** PEPut purification.

**A.** SDS-PAGE showing overexpression of the His-MBP tagged PEPut in Rosetta (DE3) *E. coli*. From left to right, lanes show protein MW markers (M), the insoluble uninduced (1), soluble uninduced (2), insoluble induced (3) and soluble induced (4) fractions. The expressed soluble His-MBP tagged PEPut is shown by a red rectangular.

**B.** SDS-PAGE showing the final purified untagged PEPut after removal of the His-MBP tag with TEV proteolysis. Lanes from left to right show protein MW markers (M) and fractions from the flow through the HisTrap column containing the pure untagged PEPut (lanes 1,2 and 3).

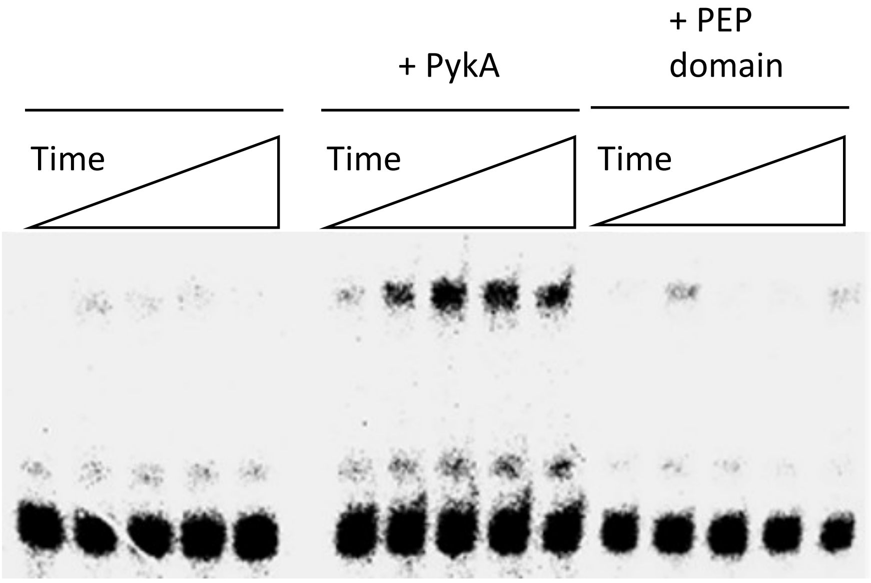


**Fig. S7:** The PEPut does not stimulate the DnaE activity.

Primer extension time course (30, 60, 90, 120 and 150 sec) assays using a primed DNA substrate (133 pM) constructed by annealing a radioactively labelled 5'-$^{32}$P 15mer primer (5'-AAGGGGGTGTGTGTG-3') onto a 110mer oligonucleotide (5'-CACACACACACACACACACACACACACACACACACACACACACACACACACACACACACCCCTTTAAAAAAAAAAAAAAAGCCAAAAGCAGTGCCAAGCTTGCATGCC-3'), at suboptimal 25 pM DnaE concentration (left), in the presence of 25 pM PykA tetramer (middle) and 25 pM PEPut domain monomer (right). At this suboptimal DnaE concentration, there is no detectable DnaE primer extension activity in the absence of PykA but clear activity is visible in the presence of PykA. By comparison, no DnaE activity is detectable in the presence of the purified PEPut domain. These data show that full length PykA stimulates the DnaE activity while the PEPut domain alone does not.

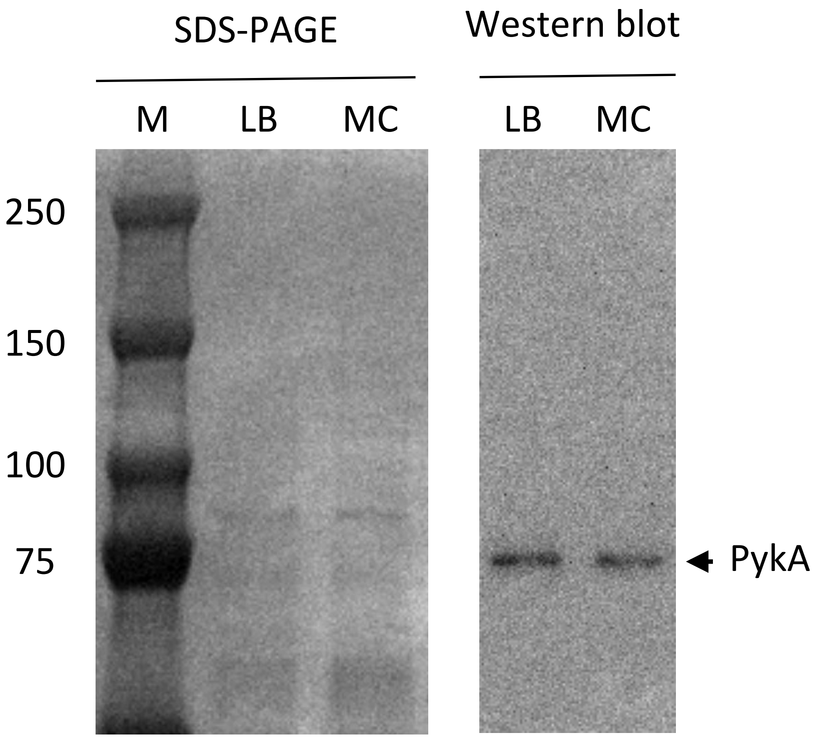


**Fig. S8:** PykA production in LB and MC medium

Cells encoding a PykA protein fused to the FLAG-tag were grown exponentially in LB and MC. At $OD_{650nm}$ = 0.5, crude extracts were prepared and 1 µg of total protein was analyzed by SDS-PAGE stained with Coomassie for total protein staining and by Western blotting. M: protein marker (kD).
